# Prior Cuing Affects Saccades to Targets in the Praying Mantis, *Sphodromantis lineola*

**DOI:** 10.1101/2023.03.30.534907

**Authors:** Théo Robert, Tan Yi Ting, Dune Ganot, Yi Jie Loh, Vivek Nityananda

## Abstract

External cues bias human attention and the perception of subsequent targets. Little is known about how cue properties, such as depth, influence insect attention. One robust cue to depth is stereoscopic disparity, the difference in the position of an object in the views of the two eyes. Praying mantises are known to use disparity to judge the distance to prey and are therefore ideal insect models to investigate its role in attention. We investigated how three cue properties- position, duration and stereoscopic disparity affect mantis selective attention towards subsequent targets. We fitted mantises with 3D glasses and presented them with a cue in 2D or in 3D, followed by two 3D stimuli: a high contrast target and a distractor at different contrasts. Our results show that cue position and distractor contrast had the most influence on responses to targets, with no strong effect of disparity. Compared to the Uncued condition, cues in two of our disparity conditions reduced target responses if presented on the opposite side of the screen, when the distractor was absent. The cues affected subsequent selective attention even when they did not themselves attract head saccades, suggesting covert but not overt attention to the cues. Our results show that the position of prior cues can affect mantis selective attention and add further evidence for the complexity of attention-like processes in insects.

**Summary Statement:** Cue position and, to a lesser extent, disparity affect praying mantises’ responses to a subsequent target. This adds further evidence for the complexity of attention-like processes in insects.

## Introduction

Attention is an essential process by which animals filter out multiple stimuli to focus on the ones of interest (Johnson and Proctor, 2004; Wolfe and Horowitz, 2004). Spatial attention, the allocation of attention to particular regions in space, has been a particularly well studied aspect of attention and the visual properties that guide it are well established, especially in humans and non-human primates (Chica et al., 2013; Krauzlis et al., 2013; Posner, 1980; Spence and Santangelo, 2009). Yet other animals, including insects, would also benefit from attentional processes – think of a bee choosing one amongst many flowers in a meadow or a mantis choosing between prey. Studies focusing on attention in insects often do so under the framework of selective attention – the ability to choose one amongst several targets (for reviews, see (De Bivort and Van Swinderen, 2016; Nityananda, 2016)). Insect selective attention is influenced by different properties of targets, including contrast. However, we do not know how prior cuing influences insect selective attention to targets presented with various properties.

A classic paradigm to study visuospatial attention in humans uses cues to bias spatial attention prior to the presentation of a target (Posner, 1980). In these studies, prior cuing influences the detection and perception of subsequent targets, depending on the position of the cues. Only a few studies have applied similar cuing paradigms in insects (Lancer et al., 2019; Sareen et al., 2011; Wiederman et al., 2017), and they have not investigated how the properties of the cue could affect insect attention. These studies also focussed on insect spatial attention within a single plane and have not investigated the influence of depth-related cues. In particular, the interaction between insect attention and stereoscopic disparity remains largely unknown. Disparity is the difference in the retinal position of an object on the two eyes. Since this difference is greater for nearer objects and lower for objects that are further away, disparity can provide information about the depth of an object independent of other visual characteristics, including angular size and motion. The ability to use disparity to calculate depth is called stereopsis, which is present in several animals (Nityananda and Read, 2017), including one group of insects, the praying mantids.

Stereopsis has been demonstrated behaviourally in mantises by measuring their probability to strike at prey presented with different disparities (Rossel, 1983). Mantises preferentially strike at prey with disparities corresponding to nearer distances - in a range typically 2.5 to 5 cm away (Maldonado et al., 1967; Nityananda et al., 2016a; Nityananda et al., 2018; Rossel, 1983). Early studies used prisms to manipulate the disparity of prey (Rossel, 1983). More recently, this was done using an insect 3D cinema, which involves affixing colour filters on each eye of the mantis, enabling the presentation of different stimuli to each eye on a computer monitor (Nityananda et al., 2016b) and thus manipulation of the perceived disparity of an object on the screen (for an example, see the schematics in Fig. 1). This method has allowed researchers to investigate the mechanisms underlying stereopsis and its role in detecting and striking at prey. Recent neurophysiological work (Rosner et al., 2019) has also suggested the possibility that disparity differences could guide attention in 3D space.

**Figure 1:**
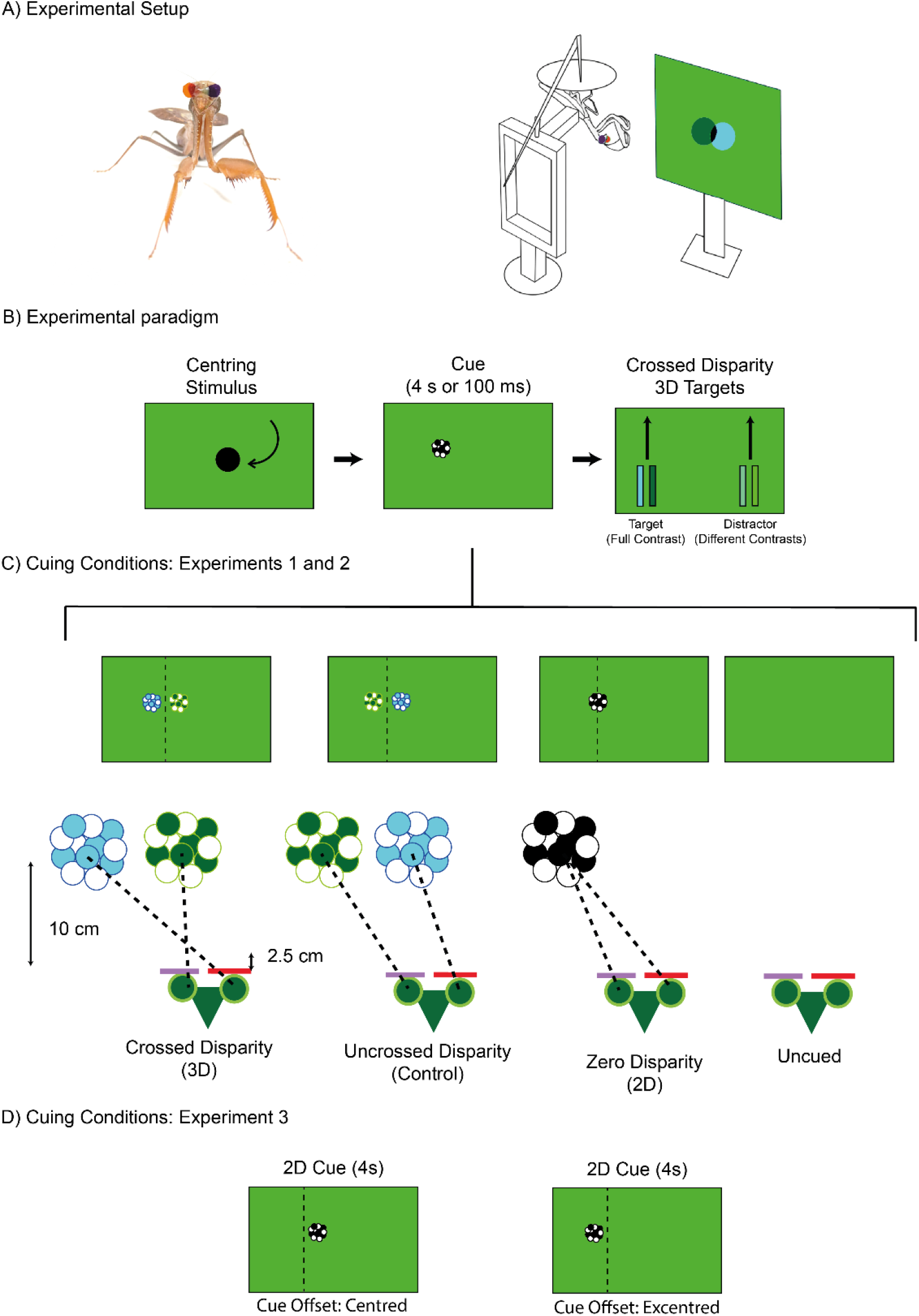
Experimental paradigm. (A) Photo of a mantis with 3D glasses attached (left). Schematic of the experimental setup representing a mantis equipped with 3D glasses and fixed upside-down in front of the computer screen on which the motivation stimulus is displayed (right). This stimulus consisted of a 3D circle spiralling inward that was used before and after each experimental block to test the motivation of the animal. (B) Schematic representation of the experimental design. We first displayed a centring stimulus spiralling towards the centre of the screen. A cue was then presented on the left or the right side of the screen with one of the four disparity conditions, either for 4 s (Experiment 1) or 100 ms (Experiment 2). After the cue disappeared, two crossed disparity rectangular stimuli were displayed, sliding upward. The target was always displayed with a contrast of 1 (full contrast) while the distractor had one of five contrasts: 0 (invisible), 0.25, 0.49, 0.75 and 1 (equal to the target). The positions of the cue, the target and distractor were counterbalanced and were presented on both the left and right sides on different trials. (C) Schematic representation of mantis’ heads wearing 3D glasses and facing the four cuing conditions in Experiments 1 and 2. The Crossed Disparity condition simulated a cue at a depth of 2.5 cm from the mantis. The Uncrossed Disparity condition had a cue with the same absolute parallax but with the positions in the right and left eye swapped, simulating an ‘impossible’ stimulus without any clear depth which served as a control condition. The Zero Disparity condition presented the cue on the screen at a depth of 10 cm from the mantis. No cue was presented in the Uncued condition. (D) Schematic representation of the cuing conditions in Experiment 3. The cue was a 4 s, zero disparity cue. This was offset either towards the centre of the screen (centred) or the edge (excentred) to match the monocular positions of the crossed and uncrossed disparity cues in the other experiments. The dotted lines are provided for comparison with the stimuli above and were not present during any of the experiments. All cues are here shown to the left of the screen but were displayed on either side on different trials.

We ran a series of behavioural experiments to investigate the effects of cuing on mantis selective attention to two subsequent stimuli that differed in contrast. Additionally, we also looked at the effect of cue disparity. We used an insect 3D cinema to present mantises with cues that varied in disparity, duration and lateral location, followed by a choice between a focal high contrast target and another identical stimulus presented at different contrasts that we called a distractor. To investigate selective attention, we tested how the properties of the cue and distractor influenced mantis saccades to the target.

## Materials and Methods

### Animals

The mantises in our experiment were all adult female Sphodromantis lineola, bred in a dedicated insect housing facility or bought as 4^th^ instar nymphs from Bugz UK. They were raised in a dedicated insect facility and kept in individual plastic containers (dimensions 19.8 by 12.5 by 15.5 cm) at a constant temperature of 25°C. When individuals did not participate in experiments, they were fed 3 times a week with an adult cricket. During experiments, mantises were fed only once a week to increase their motivation to react to stimuli (Bertsch et al., 2019; Pickard et al., 2021).

### Experimental Setup

Each mantis was fitted with 3D anaglyph glasses consisting of one red and one purple tear drop shaped transparent light filter (Fig 1A, LEE® colour filters, 135 Deep Golden Amber and 797 Purple respectively) fixed in front of the eyes of the mantises using a mixture of wax and rosin on the frons (for more details, see (Nityananda et al., 2019a)). For each mantis, the side of the red and the purple filters was determined randomly. These glasses ensured that each eye could see only one of the two channels in which we presented stimuli. We were then able to manipulate stereoscopic disparity based on the position of the stimuli in each eye (Fig 1B). An attachment was also fixed with rosin and wax to the pronota of the mantises to allow us to later attach them to the experimental stand. Mantises were allowed to recover for a day after this procedure.

On experimental days, mantises were placed hanging upside down with their feet holding a platform and fixed to the experimental stand using the attachment on their pronotum (Fig. 1A). This position is their preferred hunting position and is typical of mantises in other vision experiments (Rossel, 1983; Rossel, 1996). Our setup maintained the mantises at a fixed position of 10 cm in front of the centre of a computer screen while allowing them to freely move their head and front limbs. Experimental stimuli were displayed on the monitor (DELL U2413f LED monitor, 1920×1200 pixels resolution, 60 Hz refresh rate, 51.8 by 32.5 cm). A webcam (Kinobo USB B3 HD Webcam) placed under the screen recorded the responses of the mantises at approximately 8.35 fps.

### Experimental stimuli and design

All stimuli were generated using custom-written code with Psychtoolbox (Brainard, 1997; Kleiner et al., 2007; Pelli, 1997) (version 3.0.15) in Matlab (The Mathworks Inc.,version 2012b). Stimuli were presented in 3D by presenting each eye with stimuli in different colour channels in combination with the anaglyph filters (Fig. 1B-D). In order to balance the input of each channel through the filters, the RGB gains were adjusted to be (0 1 0) in the green channel and (0 0 0.13) in the blue channel (for more details, see (Nityananda et al., 2019a)). Stimuli in all experiments were presented against a background that consisted of 50% of the maximum output of each colour taking into account these adjustments. It was thus a mixture of blue and green that would be perceived as equally ‘grey’ for each eye.

During experiments, mantises were first presented with a centring stimulus consisting of a spiralling black disc (RGB values: 0 0 0) that swirled in from the periphery to the centre of the screen (Fig. 1B, left). After a pause of 10 seconds, mantises in conditions with a cue were presented with a cue (Fig. 1B, centre) on either the left or the right side of the screen 2.8 cm from the centre line and at height of 24.3 cm from the bottom of the screen (8.05 cm above the line of sight of the mantises). The cue was a disc (subtended angle at the viewing distance of 10 cm: 11.42°) consisting of a group of black and white dots (each of diameter 18.5 px, subtending an angle of around 2.86°). This disc was stationary but in other respects was identical to that used in previous experiments (Nityananda et al., 2019b) to elicit prey capture responses when moving. Based on pilot experiments we anticipated that a stationary cue would bias spatial attention without eliciting saccades by itself, i.e., it would lead to covert and not overt attention. In overt attention, an individual moves their eyes or head to allocate their attention in space while covert attention involves the reallocation of one’s attention without eye or head movement (see (Carrasco, 2011) for further details). The change in covert spatial attention due to the cue was tested by measuring its effect on the overt saccades to subsequent targets on either side.

Immediately after the cue disappeared, two stimuli – which we called a target and a distractor-were presented on the screen (Fig 1B, right). These consisted of two black rectangles (3.5 cm tall by 0.7 cm wide, subtending a horizontal angle of 4°) displayed 7 cm from the vertical centreline of the screen, on opposite sides. The target (RGB values: 0 0 0) always had a contrast of 1 against the background while the distractor was presented in different trials with one of five contrasts inferred from the gamma corrected RGB values passed to the screen: 0 (invisible), 0.25, 0.49, 0.75 and 1 (contrast equal to the target). When the distractor had a contrast of 1, it was identical to the target and was only nominally a distractor. Contrasts, here, are computed as per the equation usually used to compute Michelson contrasts:

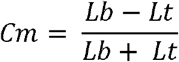

where Lt is the luminance of the target or distractor and Lb is the luminance of the background.

The target and distractor’s positions relative to the vertical centreline of the screen were shifted compared to the position of the cue to avoid a possible masking effect of the cue. Both stimuli were presented with a binocular disparity simulating a position on a virtual plane situated 2.5 cm from the mantis (Fig 1B). They appeared with their bottom edges a quarter of the screen height from the bottom of the screen (approximately 8.1 cm) and moved synchronously upward on parallel trajectories with a speed of 324 px/s (9.72 cm/s) until they reached the top of the screen. They thus took approximately two seconds after the cue disappeared to reach the vertical position of the cue. This design and movement were chosen because such stimuli (without binocular disparity) have previously been shown to elicit mantis head saccades (Rossel, 1996). In that study, when presented with two of these targets, mantises would have an equal probability to saccade towards either of them. However, if the perception of one of the targets was artificially hampered (by partially occluding it), mantises would disproportionately saccade towards the other one. As the aim of our own study was to measure how a cue can influence the perception of subsequent stimuli by the mantises through the capture of their attentional process, this paradigm appeared ideal for our experiment. Since mantises were happy to track such simple stimuli, we decided against the use of more photorealist stimuli resembling prey. These were unnecessary to test our hypotheses, could have introduced unpredictable confounding variables and made our study less comparable to older relevant work.

The presentation sides of the cue and target, and the distractor contrast were randomly assigned on a given trial and counterbalanced across trials. One experimental block consisted of two repeats of every combination of the two cue presentation sides, the two target presentation sides and the five distractor contrast values making for a total of 40 trials. A gap of 60 s was given between each trial to prevent habituation to the stimuli.

In all experiments, the motivation of the mantis was first tested by presenting a 3D black disc simulated to be on a virtual plane situated at a distance of 2.5 cm from the animal (Fig 1A, right). This disc resembled the centring stimulus (but with crossed disparity) and spiralled in from the periphery to the centre of the screen. The experiment started only after the mantis attempted to strike at the target for two consecutive presentations of this stimulus. The same stimulus was also run at the end of the experimental block and the mantises’ responses or lack of response to two consecutive presentations of the stimulus was noted. The mantis was deemed unmotivated for the experimental block if it did not track any of the stimuli in the block and also did not respond to the motivational stimulus at the end. In these cases that block was discarded. When possible, the same block was presented again to the mantis either on the same or a different day. In all 8 out of 286 experimental blocks were discarded.

We ran two experiments investigating the effect of cuing on selective attention and a control experiment designed to check whether the monocular location of the cue was responsible for the effects measured with crossed and uncrossed disparity cues.

#### Experiment 1: Testing the effect of cue position and disparity

In this experiment, the cue was presented for a duration of 4 seconds (N=14 mantises) before the stimuli appeared. The duration of the cue was similar to that previously used to study insect attention (Sareen et al., 2011).

To investigate the role of cue disparity in selective attention we had three disparity conditions (Fig. 1C, Movie 1) presented in separate experimental blocks: a Crossed Disparity condition, an Uncrossed

Disparity condition and a Zero Disparity condition. We also had an Uncued condition where no cue was presented.

In the Crossed Disparity condition, the cue disparity simulated a cue at a depth of 2.5 cm when viewed directly from 10 cm away, i.e. at the same depth as the subsequent 3D stimuli (the target and distractor). In the Uncrossed Disparity condition, the cue had the reverse disparity as in the Crossed Disparity condition, i.e. the same parallax but with the stimulus positions in each eye swapped. This therefore simulated an ‘impossible’ stimulus without any clear depth and served as a control condition. In the Zero Disparity condition the cue was displayed with images of both eyes perfectly overlapping on the screen (i.e. zero screen parallax), at a depth of 10 cm from the mantis (and a direct diagonal distance from the mantis of 10.38 cm).

These conditions were presented in a random order in a blocked design and the entire set of conditions were presented twice to 13 mantises and once to one mantis that died before the end of the experiment. Except for this last mantis, across all experimental blocks, we had eight trials per mantis for every combination of cue condition and distractor contrast value in which the cue was on the same side as the target. We also had eight trials per mantis for every combination of cue condition and distractor contrast value in which the cue was on the opposite side to the target.

#### Experiment 2: Testing the effect of cue duration

In addition to the spatial position and simulated depth of a cue, its duration could also have an impact on the ability to capture attention. To look at the effect of cue duration we therefore repeated Experiment 1 with a cue duration of 100 ms (Movie 2). We ran this experiment with 16 mantises (6 of which participated to the experiment 1).

#### Experiment 3: Controlling for the effect of the cue monocular position

In the Crossed and Uncrossed Disparity conditions of the previous experiments, the monocular position of the cue in each eye differed compared to the Zero Disparity condition. We therefore ran an experiment to test whether any difference in response between conditions was due to disparity or monocular position. Here we presented each of 13 mantises with a cue of 4 s duration in only the Zero Disparity condition but displaced 1.05 cm either toward the left or the right of its original position (Fig. 1D, Movie 3). The resultant positions of the cue corresponded to the monocular positions of the cue in the Crossed Disparity and Uncrossed Disparity condition. This manipulation offset the cue either toward the centre of the screen (hereafter “centred”) or toward the outside of the screen (hereafter “excentred”). All other details of the cue and target presentation were as for the Zero Disparity condition in Experiment 1.

### Quantification and statistical analysis

Mantis responses on every trial were recorded blind to the stimuli presented. Cue presentations were indicated on the screen by a small black disc at the bottom of the screen that was visible in the recordings but hidden from the mantis by a card barrier.

Video recordings of every trial were visually examined and the presence or absence, and the direction of mantis first head saccade were recorded. A first head saccade was defined as a distinct rapid head movement towards one side of the screen made by the mantis in the horizontal plane, allowing the mantis to place either the target or distractor in her fovea. Slow head movements were not counted and trials where the mantis started with its head already tilted toward a side of the screen were excluded from further analysis. After the video analysis, responses were matched with the location of the target and distractor. Trials in which the mantis’s first saccade was performed toward the target were recorded as 1 and trials in which the first saccade was toward the distractor or in which mantises did not make saccades were recorded as 0.

We also manually recorded the video frame on which the target and distractor appeared, the one on which the mantis started its first saccade for all trials from the experiment 1 and 2. We used these to compute the timing of the first saccade.

If the mantis performed a saccade toward the cue, before the target and distractor appeared, the trial was excluded from further analysis. In total, 2.4% of trials (229 out of 9400) were excluded from analysis due to saccades to the cue. Statistical analyses were conducted in R 4.0.4. by fitting generalised linear mixed effect models using the lme4 package (Bates et al., 2015). All models included the mantises’ ID as a random effect. We used target saccade probability as the dependent variable, coding trials with a first saccade toward the target as 1 and trials with a first saccade toward the opposite side or no saccade as 0. These models were fit using a binomial family and a logit link function.

For the analysis of the results, we combined data from Experiments 1 and 2 in a model. The independent variables in the model were the contrast of the distractor taken as a continuous variable, the cue duration, and cuing condition. Since we had two possible locations of the cue (target side cued or non-target side cued) for each of the three cuing disparity conditions and an additional category of uncued trials, there were a total of seven different cuing condition levels included in the last variable. Additionally, we also considered the interaction between the three independent variables in our analysis.

To determine the significance of an effect and select the most appropriate model, we compared the goodness of fit of models of increasing levels of complexity. The goodness of fit comparisons were conducted using the Likelihood Ratio tests from the anova() function in R. We first compared models with only one of the three main effects (distractor contrast, cue duration and cuing condition) to a null model to analyse whether any of these variables influenced the probability of first saccades towards the target. We next examined whether the effect of distractor contrast was different depending on the cuing condition by comparing a model including the main effects of distractor contrast, cueing condition and their interaction (contrast * cuing condition) with a model including only the distractor contrast). Finally, we analysed whether this last interaction was different depending on the duration of the cue. To do so, we analysed the significance of the three-way interaction by comparing the full model (contrast*cuing condition*duration) to a model including the main effects and interactions between only distractor contrast and cuing condition.

Because conditions when the distractor was not visible appeared to be of particular significance, we fitted additional models on the binomial data of the mantises’ saccades towards the target when the distractor contrast was equal to 0. We first compared a model with the cueing condition as an independent variable to the null model. We then tested the effect of the cue duration against the same null model. Finally, we tested the effect of the interaction of the cue duration with the cueing condition by comparing a model with this interaction to the model including only the effect of the cueing condition.

We also investigated whether the cue position (displayed on the side of the target or the non-target side) had an effect on the probability of mantis head saccades toward the target, specifically when the distractor was absent. To do this, we focussed on the data when the distractor contrast was 0. We excluded all Uncued trials (trials without a cue displayed) from these data since cueing position is not meaningful for the Uncued condition. We first tested the effect of the cue position and the cue disparity by comparing two models fitted with each of these variables to a null model. We then tested the interaction of the cue position with the cue disparity by comparing a model including this interaction to the model with the cue position only.

Finally, we also used the Uncued condition trials from Experiments 1 and 2 to analyse the effect of distractor contrast on the overall mantis saccade probability to either stimulus. We used a model fitted with a binomial family and a logit link function on the data from both experiments. The effect of the contrast was assessed by comparing a model with only distractor contrast as an independent variable to the corresponding null model. To confirm that an eventual effect of the distractor contrast detected by our model was not only due to the particular case when the distractor was not displayed, we also performed the same analysis on the data with the zero distractor contrast condition excluded.

To analyse the results from Experiment 3, the model selection process was similar to the previous one. First, we compared three models with each of the main effects of distractor contrast, cue position (target side or non-target side) and the offset type (centred or excentred) to a null model. Then, we compared a model with the main effects of distractor contrast and cue position with a model that also included their interaction. Finally, we compared this last model to a full model including the offset side.

To analyse the timing of the first saccade, we used mixed effect models fitted on the number of video frames taken by the mantises to perform their first saccades after the target and distractor appeared on screen. To do so, we use the glmmTMB package (Brooks et al., 2017) to fit models with a negative binomial family and a log link function, and also included the modelling of the heteroscedasticity by including a dispersion formula based on the interaction between the cue disparity and the cue duration. All models included the mantis identity as random effect. With these models we examined the effect of the distractor contrast, the cue duration, cue position (target side or non-target side), cue condition (Crossed, Uncrossed, Zero disparity or Uncued) as well as side of the first saccade (towards the target or toward the distractor) by comparing the fit of model with each of these main effect to a null model. We then built a model with all significant main effects and progressively added interaction effects and compared each new model to the one preceding it.

For every analysis, after our model comparison, we examined the significance of the estimates of the best model using Wald tests through the summary() function in R. In addition, we calculated the 95% confidence intervals of these intercepts to confirm these test results (see supplementary information).

We used a significance level of α=0.05 throughout our analyses.

## Results

The best model of the target saccades in Experiments 1 and 2 showed that there was a significant main effect of the distractor contrast (χ^2^=697.570, df=1, p<0.001) and a significant interaction effect of distractor contrast and cuing condition on saccade probability (χ^2^ =29.287, df=12, p=0.004) but no significant effect of cue duration (χ^2^ =23.205, df=14, p=0.057).

### Higher contrast distractors reduce target saccade probability

The probability that the first saccade was to the target decreased significantly as distractor contrast increased (Fig. 2A and B, Table S1, Estimate ± standard error=-2.894±0.205, Z=-14.096, p<0.001). This effect can likely be explained by the fact that, across conditions, as the distractor became more discernible from the background, mantises were more likely to saccade to the distractor, and therefore the probability that they saccaded toward the target decreased (for more details and supporting statistics, see Fig. S1).

**Figure 2:**
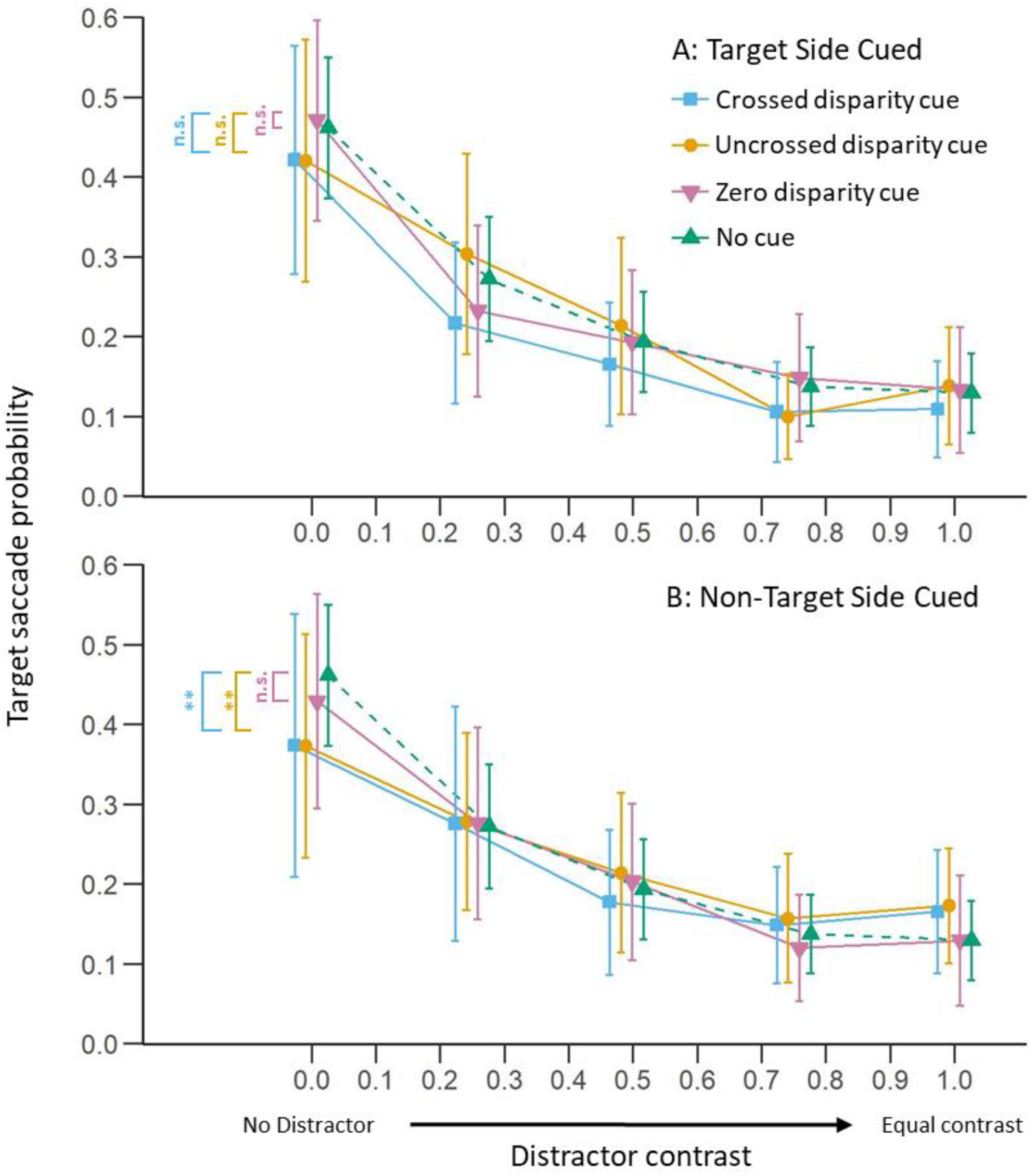
The influence of cue position and disparity on target saccade probability. Mean (± 95% C.I.) proportion of trials with a first saccade towards the target as a function of the contrast of the distractor (N=24 mantises). The target always had contrast of 1. The cue was presented either on the side of the target (A) or on the side opposite to the target (B). Blue curves represent trials with crossed disparity cues, yellow curves are for trials with uncrossed disparity cues while pink curves are for trial conducted with zero disparity cues. Green curves represent uncued trials and are the same across the two panels. The comparisons to the left are all with respect to the Uncued condition. Both the intercepts and the slopes of the Crossed and Uncrossed Disparity conditions were significantly different from the Uncued conditions in the panel B. When comparing between the disparity conditions (not shown), only the slopes of the Uncrossed and Zero Disparity conditions in B were significantly different (see text for further details). Results combine data from both 4 s (Experiment 1) and 100 ms (Experiment 2) cues. ** denotes a p-value < 0.01, n.s. denotes no significant differences.

The effect could further be explained by a second phenomenon: the overall likelihood that the mantises performed any saccade at all, to either stimulus, also decreased as the target and distractor had more similar contrasts (Fig. 3 ; Effect of the distractor contrast: Estimate ± standard error=-1.544±0.159, Z=-9.690, p<0.001). The curve presented in figure 3 shows that the decrease in saccades is particularly pronounced between the 0 and 0.25 distractor contrasts. This indicates that the effect of the contrast shown in our model is mostly driven by the presence of the distractor. However, the effect of distractor contrast was significant even if we excluded the condition without a distractor. This suggests that it is the distractor contrast and not merely the presence of distractor that reduces the saccades to the target (Estimate ± standard error=-0.513±0.230, Z=-2.233, p=0.026). The presence of the distractor thus strongly reduced the overall number of saccades and the increase in distractor contrast reduced the saccades to the target.

**Figure 3:**
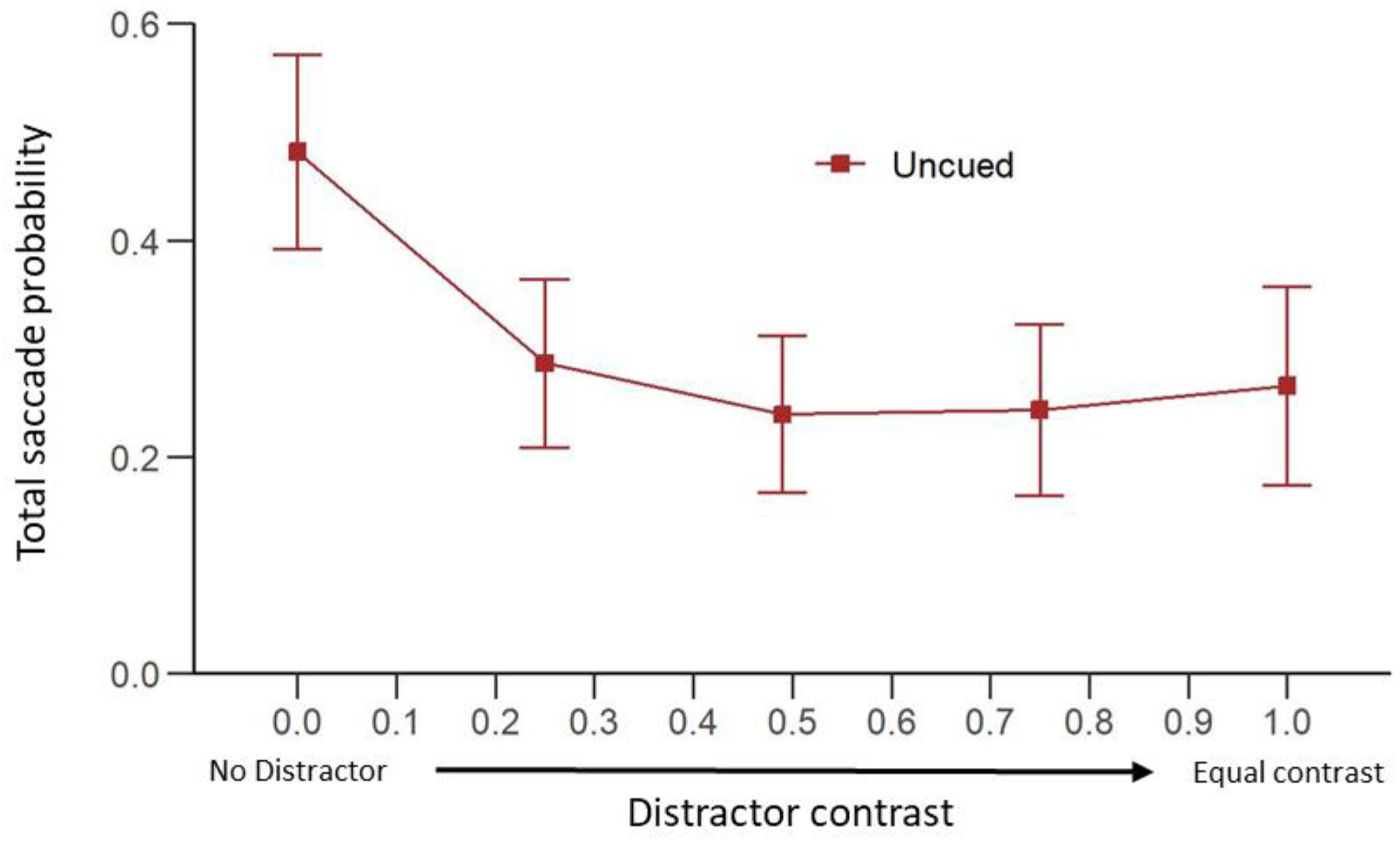
Distractor contrast affects overall saccade probability. Mean (± 95% C.I.) of the individual proportion of Uncued trials with at least one saccade towards either the target or the distractor as function of distractor contrast (N=24 mantises).

### The effect of cue position and disparity on target saccade probability

When the cue was presented on the side of the target, it did not significantly affect target saccade probability compared to the Uncued condition (Fig. 2A, Table S1, Crossed vs Uncued condition: Estimate ± standard error=-0.268±0.157, Z=-1.710, p=0.087 ; Uncrossed vs Uncued condition: Estimate ± standard error=-0.189±0.156, Z=-1.213, p=0.225 ; Zero Disparity vs Uncued condition: Estimate ± standard error=-0.189±0.156, Z=-1.211, p=0.226). In these cases, there was also no significant interaction effect between distractor contrast and the cue type on target saccade probability compared to the Uncued condition (Crossed vs Uncued condition: Estimate ± standard error=-0.006±0.315, Z=-0.018, p=0.985 ; Uncrossed vs Uncued condition: Estimate ± standard error=0.359±0.300, Z=1.197, p=0.231 ; Zero Disparity vs Uncued condition: Estimate ± standard error=0.334±0.302, Z=1.106, p=0.269).

However, when the cue was presented on the opposite side to the target, it had different effects for different distractor contrasts. When the distractor contrast was zero, only the target was visible. In this case, compared to the Uncued condition, the crossed and uncrossed disparity cues reduced the target saccade probability (Fig. 2B, Fig. 4D & E, Table S1, Crossed vs Uncued condition: Estimate ± standard error=-0.434±0.157, Z=-2.765, p=0.007 ; Uncrossed vs Uncued condition: Estimate ± standard error=-0.441±0.156, Z=-2.830, p=0.005). This was not true of the Zero Disparity condition (Fig. 4F, Zero Disparity vs Uncued condition: Estimate ± standard error=-0.195±0.156, Z=-1.251, p=0.211) where the cue was presented 10 cm away from the mantis. The effect sizes for crossed and uncrossed disparity cues were correspondingly larger than for zero disparity cues. This suggests that in this situation, cues in two of the three disparity conditions covertly attracted the mantises’ attention compared to a condition without a cue, making them less likely to look towards a target subsequently presented on the other side of the screen. Importantly, the cues themselves did not result in saccades, since we had excluded trials where this happened. This result is therefore not explained by an initial change in head direction before the saccade to a target.

**Figure 4:**
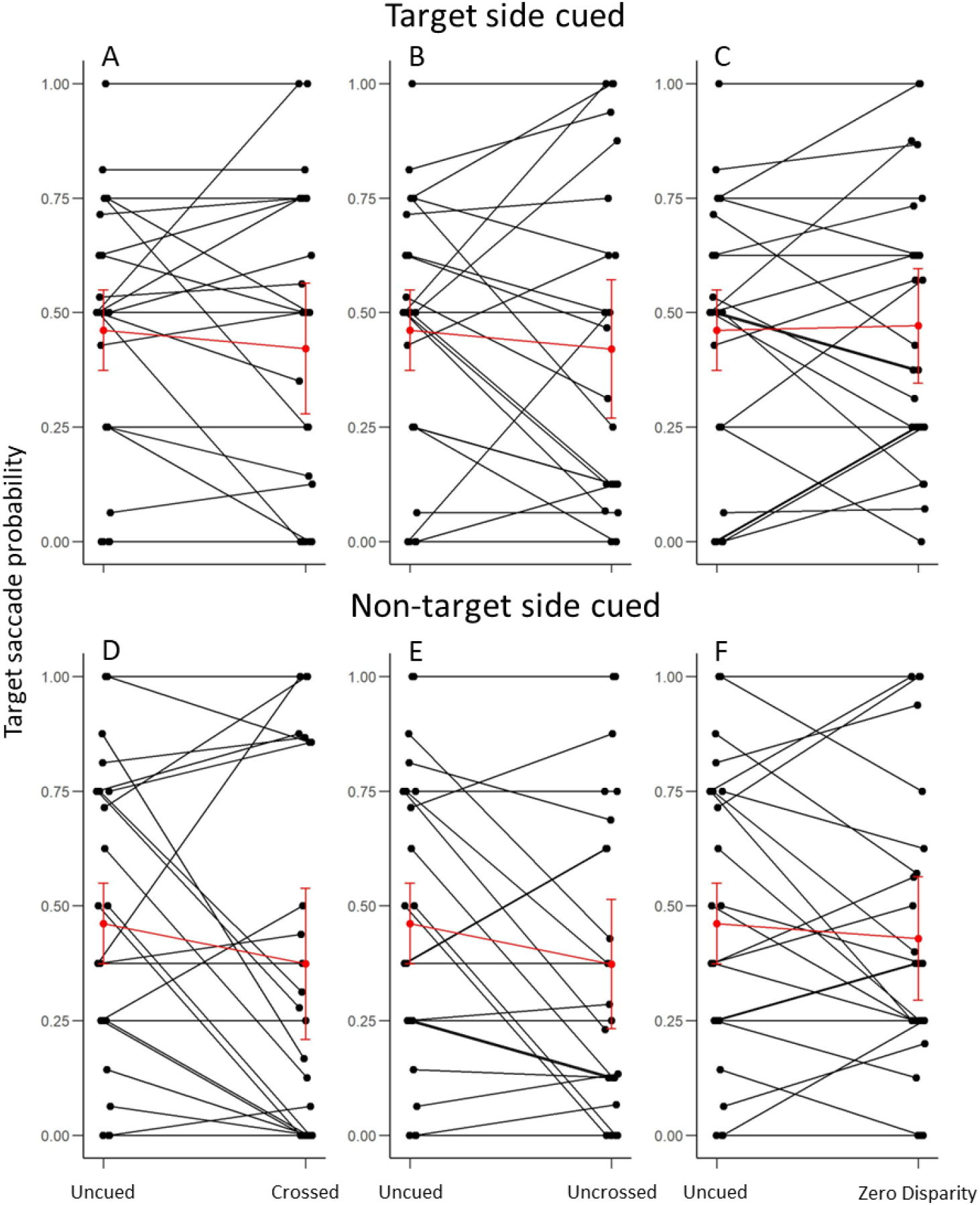
Cuing conditions effects on individual target saccade probability in the absence of a distractor. Data are shown for when the cue was on the same side as the target (top row) or the opposite side (bottom row). The left points of each graph represent the target saccade probabilities in the Uncued condition. The right points represent the individual probabilities for the Crossed (A, D), Uncrossed (B, E) and the Zero Disparity condition (C,F) Black points and lines represents data from individual mantises. Red points and lines show the average probabilities (± 95% C.I.) for all the mantises (N=24).

The decrease in target saccade probability with increasing distractor contrast was significantly attenuated when these cues were on the non-target side (Table S1, Crossed vs Uncued condition: Estimate ± standard error=0.887±0.295, Z=3.011, p=0.003 ; Uncrossed vs Uncued condition: Estimate ± standard error=1.112±0.288, Z=3.857, p<0.001). As the contrast of the distractor increased, the crossed and uncrossed disparity cues thus appeared to progressively have a reduced effect, and the target saccade probability became comparable to that in the Uncued condition (Fig. 2B). This significant interaction effect between cue and contrast was, however not seen for a Zero Disparity cue (Zero disparity vs Uncued condition: Estimate ± standard error=0.389±0.300, Z=1.300, p=0.194).

Despite the different effect of cues with different disparities compared to the Uncued condition, we did not find many differences when comparing the effects of these cues with each other. When presented on the non-target side, the main effect of a zero disparity cue was not significantly different to that of both crossed and uncrossed disparity cues (Table S2, Crossed disparity vs Zero Disparity condition: Estimate ± standard error=-0.239±0.182, Z=-1.318, p=0.188 ; Uncrossed disparity vs Zero Disparity condition: Estimate ± standard error=-0.247±0.181, Z=-1.366, p=0.172). The interaction with contrast also did not significantly differ for zero disparity and crossed disparity cues (Estimate ± standard error=0.498±0.339, Z=1.467, p=0.142). However, the effect of this interaction with contrast did significantly differ between the zero disparity and uncrossed disparity conditions (Estimate ± standard error=0.722±0.334, Z=2.165, p=0.030). Taken together, these results show that disparity only seems to have a limited influence on target saccade probability.

To confirm the results for this previous model, we focused our analysis on the trials when the distractor was not visible (distractor contrast=0). In this specific case, the model selection phase confirmed the effect of the cuing condition (Fig. 4; χ^2^ =13.958, df=6, p=0.030). The examination of the estimates of this particular model including the effect of the cueing condition also confirmed that when the cue was on the side opposite to the target with a Crossed Disparity (Table S3, Estimate ± standard error=-0.575±0.200, Z=-2.869, p=0.004) or an Uncrossed Disparity (Estimate ± standard error=-0.556±0.200, Z=-2.775, p=0.006), it decreased the probability of head saccades towards the target compared to the Uncued condition. However, when compared to a condition with a Zero Disparity cue in the same position, the effect of the Crossed (Table S4, Estimate ± standard error=-0.318±0.232, Z=-1.375, p=0.169) and Uncrossed Disparity cues (Estimate ± standard error=-0.299±0.231, Z=-1.293, p=0.196) did not significantly differ.

Crossed and Uncrossed Disparity cues appear to have an effect on mantis saccades to the target but only when the cues were on the non-target side and when the distractor was not visible. We therefore decided to analyse the effect of the cue position (on the side of the target or the opposite side) for trials when the distractor was not shown. For these models, we used the data of the trials with a distractor contrast of 0 and excluded the Uncued condition because the cue position was not relevant for this condition. The model selection showed that the cue position had a significant effect (χ^2^ =3.851, df=1, p=0.0497). However, in this specific condition, cue position did not interact with cue disparity (χ^2^ =4.450, df=4, p=0.349). Thus, the selected model indicated that the chances of saccades toward the target were lower when the cue was on the opposite side of the screen than when it was on the side of the target (Table S5, Estimate ± standard error=-0.269±0.136, Z=-1.975, p=0.048).

Overall, cues seemed to attract attention away from targets when on the opposite side but not boost attention when on the same side. In addition, the disparity of the cue only had a limited effect.

### Monocular cue position of a zero disparity cue does not affect target saccade probability

The monocular position of stimuli differs for zero disparity cues and the crossed and uncrossed disparity cues (Fig. 1C). We therefore ran an additional experiment to verify whether any differences between cuing conditions observed with the crossed and uncrossed disparity cues was due to the disparity or the monocular position of the cue. We presented a zero disparity cue offset at either of the monocular positions of the cues with disparity used above. Similar to the results above, target saccade probability significantly decreased with an increase in distractor contrast (Fig. 5; χ^2^ =52.920, df=1, p<0.001; Effect of distractor contrast: Estimate ± standard error=-1.273±0.179, Z=-7.126, p<0.001). However, the position of the monocular cue did not significantly decrease target saccade probability (χ^2^ =1.795, df=1, p=0.180). There was also no significant effect of the direction of the cue offset (χ^2^ =0.210, df=1, p=0.647). The offset did not have a significant interaction with distractor contrast (χ^2^ =0.859, df=2, p=0.651). Finally, the three-way interaction between distractor contrast, cue position and offset direction was also not significant (χ^2^ =5.819, df=6, p=0.444). Overall, these results confirm that the monocular position of a zero disparity cue does not have an effect on target saccade probability.

**Figure 5:**
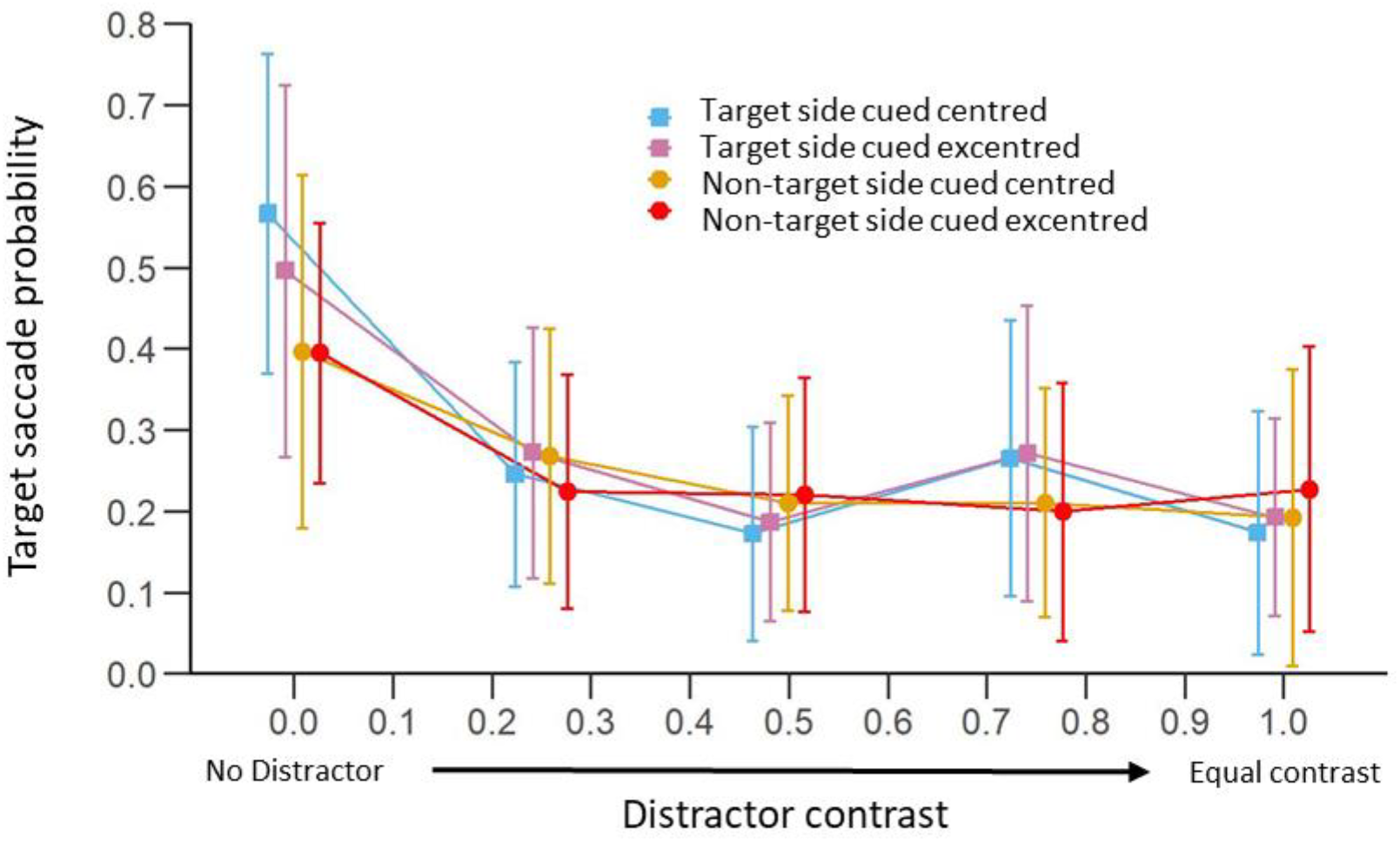
Monocular cue position of a zero disparity cue does not affect target saccade probability. Mean (± 95% C.I.) proportion of trials with a first saccade towards the high contrast target as a function of distractor contrast (Experiment 3, N=13 mantises). The cue had a duration of 4 s and was either shifted towards the centre of the screen (centred) or towards the outside (excentred) of the screen to match the monocular positions of the cue in the Crossed Disparity condition. Blue and purple curves show results for trials with the cue on the side of the high contrast target and shifted toward the centre and towards the outside of the screen respectively. The yellow and red curves represent the trials with the cue on the opposite side and shifted towards the centre and towards the outside of the screen respectively.

### Latency of first saccade is influenced by distractor contrast and the presence of the cue

Mantises took less time to perform their first saccade when it was directed at the target than when it was towards the distractor (Fig. 6; Effect of saccade direction: χ^2^ =49.976, df=1, p<0.001 ; Estimate ± standard error=-0.285±0.050, Z=-5.720, p<0.001). However, a significant interaction with the distractor contrast showed that while the latency to perform a saccade towards the target increased with the distractor contrast (Effect of distractor contrast: χ^2^ =133.510, df=1, p<0.001 ; Estimate ± standard error=0.293±0.027, Z=10.780, p<0.001), this increase was not seen for saccades towards the distractor (Interaction: χ^2^ =27.143 ; df=1, p<0.001 ; Estimate ± standard error=-0.345±0.066, Z=-5.21, p=0.395).

**Figure 6:**
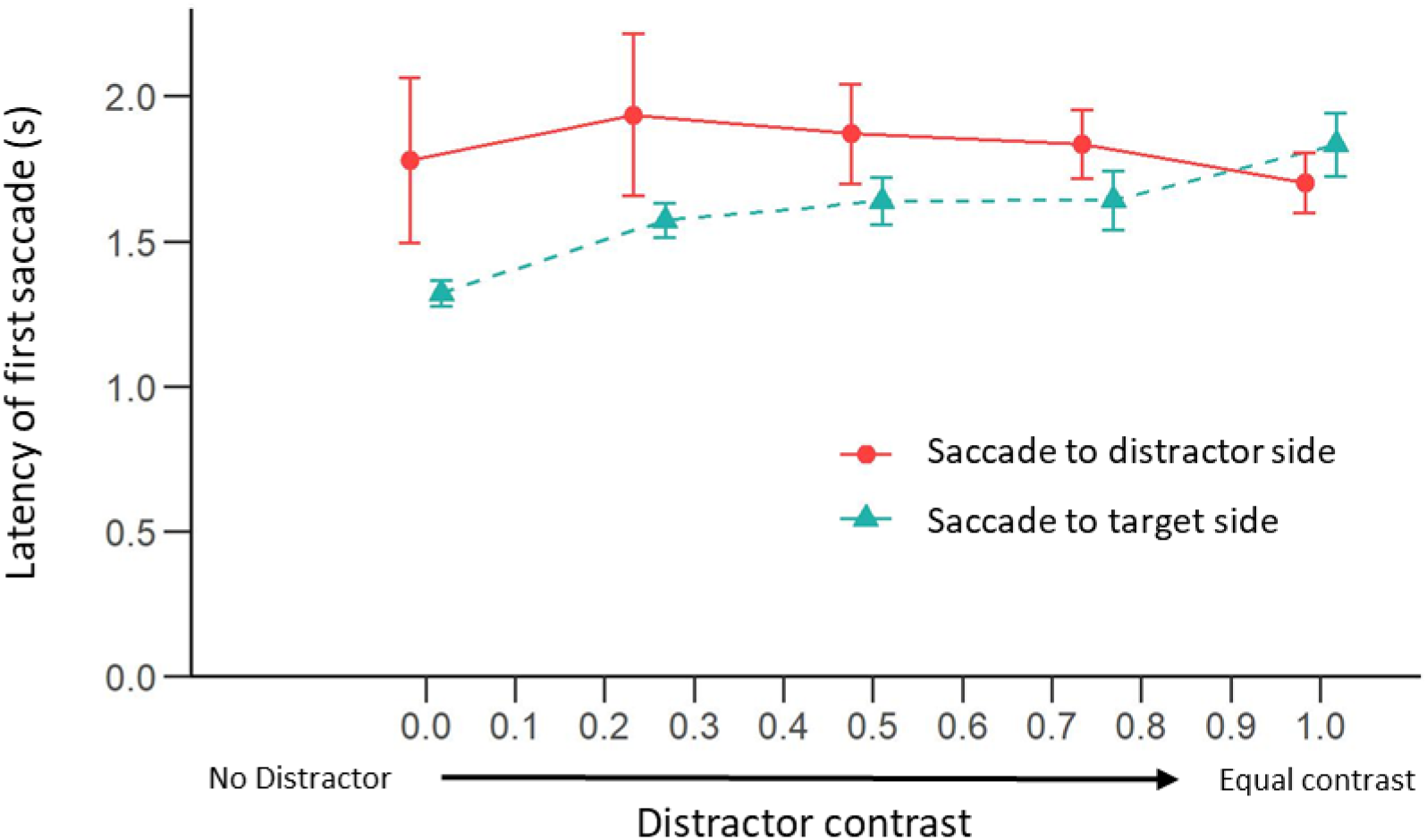
The latency to perform the first saccade increases with the distractor contrast when the saccade is towards the target but not when it is towards the distractor. Mean (± 95% C.I.) latency taken by the mantises to perform their first saccade after the target and distractor appeared as a function of the distractor contrast (Experiment 1 and 2, N=24 mantises, N=2685 saccades). The turquoise curve shows the latency of saccades towards the target and the red curve shows the latency of targets towards the distractor.

In addition, the presence of the cue also affected the latency to perform the first saccade. Mantises took more time to perform their first saccade in trials with a cue of any disparity condition compared to the Uncued condition (Fig. 7; Effect of cue type: χ^2^ =14.460 ; df=3, p=0.002 ; Crossed disparity cue vs Uncued: Estimate ± standard error=0.057±0.024, Z=2.330, p=0.020 ; Uncrossed disparity cue vs Uncued: Estimate ± standard error=0.066±0.024, Z=2.760, p=0.006 ; Zero disparity cue vs Uncued: Estimate ± standard error=0.072±0.023, Z=3.080, p=0.002). However, there was no difference between the different cue conditions (Crossed vs Uncrossed disparity cue: Estimate ± standard error=0.010±0.023, Z=0.420, p=0.674 ; Crossed vs Zero disparity cue: Estimate ± standard error=0.016±0.022, Z=0.700, p=0.485 ; Uncrossed vs Zero disparity cue: Estimate ± standard error=0.006±0.022, Z=0.270, p=0.784). Moreover, there was no interaction with the distractor contrast and the saccade direction (Effect of interaction: χ^2^ =12.674, df=9, p=0.178).

**Figure 7:**
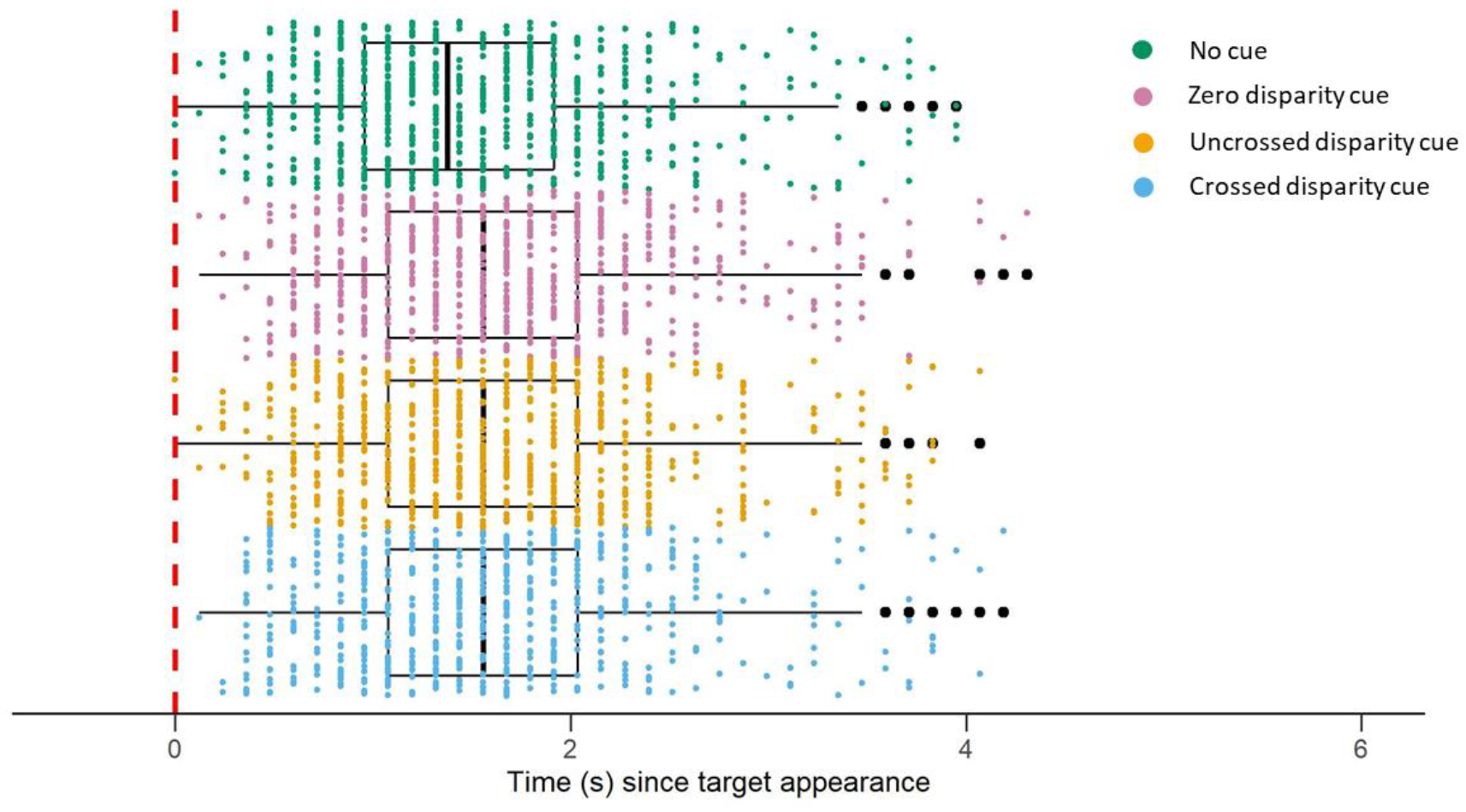
Cue presence increases the latency to perform the first saccade. Boxplots showing the distribution of latencies to perform the first saccade depending on the cue type. Dots represent individual saccades performed in trials with crossed disparity cues (blue), uncrossed disparity cues (yellow), zero disparity cues (pink) or uncued conditions (green). The red dashed line represents the time of appearance of the target and distractor.

Finally, the cue duration or saccade side (on the side of the target or of the distractor) did not have an effect on the timing of mantis first saccades (Effect of cue duration: χ^2^ =0.032 ; df=1, p=0.858 ; Effect of cue side: χ^2^ =0.126 ; df=1, p=0.723).

## Discussion

Our results showed that prior cuing influences selective attention in mantises. The presence of a cue, without any overt behavioural response, nonetheless influenced subsequent selective attention between two stimuli. Mantis selective attention was influenced mostly by the contrast of the distractor and cue position and was not strongly affected by the disparity of the cue.

### Mantis attention to simultaneous stimuli

As both our stimuli grew more identical, mantises struggled to selectively choose either of the stimuli. This might reflect poor selective attention or divided attention to both stimuli. Mantis saccade rates have previously been shown to be reduced when more than one target or distractor was shown simultaneously in a trial (Yamawaki, 2006), which resembles our results. Different neural units may be responsible for triggering saccades towards different areas of mantises’ visual field (Yamawaki, 2006). When several units are activated at the same time, they would compete to initiate a saccade, and this competition would decrease the chances of saccade. How might a mantis catch prey given this apparent limitation to their ability to choose between stimuli? The presence of two identical equidistant prey would likely be extremely rare in their natural environment. Natural differences in distance or prey traits could resolve the problem for the mantis. The selective attention mechanism of the mantis would thus appear to depend on the natural variation in the environment to enable prey selection. This points towards the importance of considering the role of ecology when studying the cognitive abilities of animals.

Other work (Rossel, 1996) presenting mantises with identical zero disparity stimuli found that their first saccades were not made clearly towards one or the other target but were more evenly distributed, including towards the centre between the stimuli. It is not clear there whether the total proportion of trials with saccades also declined in these trials with two identical stimuli compared to trials with a single target as we observed in our experiment. Flies have also been shown to choose between two stimuli as long as they are far enough apart. With stimuli that are closer together they choose the midpoint of two stimuli (Reichardt and Poggio, 1973; Reichardt and Poggio, 1975). On the other hand, in dragonflies, which are predatory insects like mantises, an interneuron ensures that one of two paired targets can be selected (Lancer et al., 2019; Wiederman and O’Carroll, 2012).

The presence of multiple stimuli has also been shown to slow down mantis reaction time in some experimental conditions (see part 3.3.2 in (Yamawaki, 2006)), similar to what we find in our results. This effect of the simultaneous presence of a distractor on the hemifield opposite to the target resembles the Remote Distractor Effect observed in humans (Walker et al., 1995; Walker et al., 1997) where distractors slow down the execution of eyes saccades to targets. However, contrary to our results, this effect in humans does not seem to be affected by the distractor contrast (Born and Kerzel, 2008).

### Effects of cue position

Cue position influenced mantis saccades in our experiment. In primates, cue position also changes the attentional performance based on the contrast (Carrasco et al., 2000; Herrmann et al., 2010; Snowden et al., 2001). However, the presence of two variable contrast targets or of a single target in these primate studies means that our results may not be directly comparable. Importantly, our cues had an effect only when presented on the side opposite to the target, when they reduce saccades to the target (Fig. 2, Fig. 4). This supported our expectation that target saccades would be lower when the cue was on the distractor side. However, we did not see enhanced attention (which would be indicated in our experiment by an increase in target saccades) when the cue was on the same side as the target, which is often seen in similar experiments with primates (Carrasco et al., 2000; Herrmann et al., 2010). Previous research on insect attention has found cuing effects that enhanced attention, but these experiments typically used cues that were identical to the targets (Lancer et al., 2022; Sareen et al., 2011). In our experiments the cue and the target were very different, and this might explain why we do not see the cue boost attention in our experiments.

Cues presented on the opposite side of a target did reduce attention to the target. This arguably resembles the effect of the presence of the distractor. However, the cue vanished before the appearance of the target, while in trials with a distractor, the target and distractor were presented simultaneously. The cue thus had an effect that persisted after it disappeared for at least a limited duration (approximately two seconds before the target reached the same vertical position). If the cue is impairing selective attention like the distractor does, this effect appears to last longer than the immediate presence of a stimulus. This was similar to previous work showing that a cue can influence fruit flies’ target selection both immediately after cuing or after a delay of 2 seconds (Sareen et al., 2011).

### Effects of cue disparity

Most studies of spatial attention in humans and non-human primates have investigated attention using cues and stimuli in a single plane. Fewer studies have investigated how attention is allocated in three-dimensional space. Studies that have researched this have found that attention can be guided by cues to depth, especially stereoscopic cues (Andersen, 1990; Andersen and Kramer, 1993; Atchley and Kramer, 1997; He and Nakayama, 1995; Plewan and Rinkenauer, 2019; Plewan and Rinkenauer, 2021; Zou et al., 2022). In our experiments, although the presence of a cue slowed down the reaction time of mantises, its disparity did not affect mantises’ latency to perform a saccade (Fig. 7). This suggests that the decision to saccade is influenced by the presence of a prior cue but not the cue properties. It is also possible that disparity or other cue properties had a stronger effect on saccade dynamics and the fine-grained details of mantis head movements. Unfortunately, our recording setup was not designed to conduct such fine analyses as our videos were stored at 8.35 frames per second. Therefore, future studies analysing mantis behaviours in response to various cuing paradigms with a finer temporal resolution would be important to investigate these effects.

Cue disparity also had little effect on mantis saccade probability in our experiments. Crossed disparity cues simulated a cue nearer to the mantis while zero disparity cues simulated a farther cue. The fact that “near” cues had a larger effect compared to “farther” cues suggest that mantis attention could be influenced more by nearer objects compared to farther ones. This would be similar to some results found in humans (Chen et al., 2012; Finlayson and Grove, 2015; Plewan and Rinkenauer, 2017). It would also seem to fit in with results showing that mantises prefer to strike at stimuli placed 2.5 cm from them and that their probability of striking is dramatically reduced for stimuli farther than 5.63 cm (Nityananda et al., 2016a). However, cues with uncrossed disparity had a similar effect to crossed disparity cues. This similarity in response to cues of crossed and uncrossed disparities is different from previous results about prey capture behaviour. There, mantises were seen to be more likely to strike at crossed disparity targets and not to uncrossed or zero disparity targets (Nityananda et al., 2016b; Nityananda et al., 2018). The difference in results suggest two points. First, mantises could attend to targets based on the parallax between monocular positions rather than target disparity. Secondly, the decision to capture prey is governed by a separate, possibly later module which does differentiate between crossed and uncrossed disparities. Previous work (Nityananda et al., 2019a; Nityananda et al., 2019b) also has argued that prey detection and prey capture are governed by separate mechanisms and our results add further support for these mechanisms. Work there has also shown that both crossed and uncrossed cue disparities can prime subsequent strikes to targets, and crossed cue disparities had a stronger effect (Nityananda et al., 2019b).

Our results can be understood in light of the neural mechanisms underlying stereopsis in mantises. Neurons tuned to specific crossed disparity signals have been recently found in the optic lobes of praying mantises (Rosner et al., 2019; Rosner et al., 2020). These are binocular neurons that connect to the optic lobes of both eyes and then feed into more central regions of the brain. More relevant to our experiments are neurons that relay information from the central complex to the optic lobe. These neurons have been suggested to serve an attentional function and could be involved in the responses we see to cues. Notably, these centrifugal neurons included neurons that responded to both stimuli simulating nearby targets (TMEcen-neurons) and stimuli with diverging lines of sight (TAcen-neurons) as would occur for our cues with crossed and uncrossed disparity respectively. The neurons that responded to diverging lines of sight responded more strongly to bright stimuli but given that our cue had both white and black dots, they still could have been stimulated in our study. Our results thus add support to the idea that these neurons could be involved in attentional processing. They further suggest that some of these neurons should potentially govern attention based on the parallax between monocular positions, or the presence of any stimulus, rather than only crossed disparities. Other neurons or a subset of these neurons would presumably then respond only to the crossed disparities and send commands to the motor circuits to enable prey capture.

The involvement of central brain structures in mantis attentional processes is also consistent with the few previous studies on insect attention. Most of these studies have shown that some of the neurons involved are present early on in the nervous system, close to the peripheral sensory neurons (Kim et al., 2015; Lancer et al., 2019; Paulk et al., 2014; Pollack, 1988; Römer and Krusch, 2000; Tang and Juusola, 2010; van Swinderen, 2012; Wiederman and O’Carroll, 2012; Wiederman et al., 2017). However, several different brain areas, including more central areas such as the mushroom bodies (Xi et al., 2008; Zhang et al., 2007) and the central complex (Seelig and Jayaraman, 2015) have also been implicated in insect selective attention.

Our results demonstrate that prior cuing can affect mantis selective attention and that these effects are compatible with the known visual neurophysiology of the praying mantis. They thus further our understanding of the sophistication of attention-like abilities in insects.

