## Supplementary Information, Tables and Figures for "Prior Cuing Affects Saccades to Targets in the Praying Mantis, *Sphodromantis lineola*"

**Table S1: Summary of effects of distractor contrast and cuing condition with the Uncued condition as baseline. A)** Estimates, standard errors, Wald test results and p-values of factors in the best model for the analysis of experiment 1 and 2. **B)** Bootstrapped 95% confidence intervals of the estimates presented in table A).

| **A)** | **Estimate** | **Std Error** | **Z Value** | **P-value** | |
|---|---|---|---|---|---|
| Intercept - Uncued | -0.454 | 0.307 | -1.482 | 0.138 | |
| Cuing Condition - Crossed Disparity Target Cued | -0.268 | 0.157 | -1.710 | 0.087 | |
| Cuing Condition - Crossed Disparity Distractor Cued | -0.434 | 0.157 | -2.768 | 0.006 | ** |
| Cuing Condition - Uncrossed Disparity Target Cued | -0.189 | 0.156 | -1.213 | 0.225 | |
| Cuing Condition - Uncrossed Disparity Distractor Cued | -0.442 | 0.156 | -2.834 | 0.005 | ** |
| Cuing Condition - Zero Disparity Target Cued | -0.189 | 0.156 | -1.211 | 0.226 | |
| Cuing Condition - Zero Disparity Distractor Cued | -0.195 | 0.156 | -1.251 | 0.211 | |
| Contrast - Uncued | -2.601 | 0.177 | -14.708 | <0.001 | *** |
| Contrast : Cuing Condition - Crossed Disparity Target Cued | -0.006 | 0.315 | -0.018 | 0.985 | |
| Contrast : Cuing Condition - Crossed Disparity Distractor Cued | 0.887 | 0.295 | 3.008 | 0.003 | ** |
| Contrast : Cuing Condition - Uncrossed Disparity Target Cued | 0.359 | 0.300 | 1.197 | 0.231 | |
| Contrast : Cuing Condition - Uncrossed Disparity Distractor Cued | 1.112 | 0.288 | 3.855 | <0.001 | *** |
| Contrast : Cuing Condition - Zero Disparity Target Cued | 0.334 | 0.302 | 1.106 | 0.269 | |
| Contrast : Cuing Condition - Zero Disparity Distractor Cued | 0.389 | 0.300 | 1.300 | 0.194 | |

| **B)** | **Estimates** | **2.5%** | **97.5%** |
|---|---|---|---|
| Intercept - Uncued | -0.454 | -1.052 | 0.119 |
| Cuing Condition - Crossed Disparity Target Cued | -0.268 | -0.591 | 0.019 |
| Cuing Condition - Crossed Disparity Distractor Cued | -0.434 | -0.769 | -0.129 |
| Cuing Condition - Uncrossed Disparity Target Cued | -0.189 | -0.480 | 0.095 |
| Cuing Condition - Uncrossed Disparity Distractor Cued | -0.442 | -0.761 | -0.140 |
| Cuing Condition - Zero Disparity Target Cued | -0.189 | -0.501 | 0.084 |
| Cuing Condition - Zero Disparity Distractor Cued | -0.195 | -0.509 | 0.114 |
| Contrast - Uncued | -2.601 | -2.978 | -2.266 |
| Contrast : Cuing Condition - Crossed Disparity Target Cued | -0.006 | -0.638 | 0.605 |
| Contrast : Cuing Condition - Crossed Disparity Distractor Cued | 0.887 | 0.265 | 1.487 |
| Contrast : Cuing Condition - Uncrossed Disparity Target Cued | 0.359 | -0.240 | 0.966 |
| Contrast : Cuing Condition - Uncrossed Disparity Distractor Cued | 1.112 | 0.525 | 1.742 |
| Contrast : Cuing Condition - Zero Disparity Target Cued | 0.334 | -0.244 | 0.937 |
| Contrast : Cuing Condition - Zero Disparity Distractor Cued | 0.389 | -0.207 | 0.974 |

**Table S2: Summary of effects of distractor contrast and cuing condition with the Zero Disparity, Distractor Cued condition as baseline. A)** Estimates, standard errors, Wald test results and p-values of factors in the best model for the analysis of experiment 1 and 2**. B)** Bootstrapped 95% confidence intervals of the estimates presented in table A).

| **A)** | **Estimate** | **Std Error** | **Z Value** | **P-value** | |
|---|---|---|---|---|---|
| Intercept - Zero Disparity Distractor Cued | -0.650 | 0.320 | -2.029 | 0.043 | * |
| Cuing Condition - Crossed Disparity Target Cued | -0.073 | 0.181 | -0.402 | 0.688 | |
| Cuing Condition - Crossed Disparity Distractor Cued | -0.239 | 0.182 | -1.318 | 0.188 | |
| Cuing Condition - Uncrossed Disparity Target Cued | 0.007 | 0.180 | 0.036 | 0.971 | |
| Cuing Condition - Uncrossed Disparity Distractor Cued | -0.247 | 0.181 | -1.366 | 0.172 | |
| Cuing Condition - Zero Disparity Target Cued | 0.006 | 0.181 | 0.032 | 0.975 | |
| Cuing Condition - Uncued | 0.195 | 0.156 | 1.253 | 0.210 | |
| Contrast - Zero Disparity Distractor Cued | -2.212 | 0.244 | -9.074 | <0.001 | *** |
| Contrast : Cuing Condition - Crossed Disparity Target Cued | -0.395 | 0.357 | -1.106 | 0.269 | |
| Contrast : Cuing Condition - Crossed Disparity Distractor Cued | 0.498 | 0.339 | 1.467 | 0.142 | |
| Contrast : Cuing Condition - Uncrossed Disparity Target Cued | -0.031 | 0.343 | -0.089 | 0.929 | |
| Contrast : Cuing Condition - Uncrossed Disparity Distractor Cued | 0.722 | 0.334 | 2.165 | 0.030 | * |
| Contrast : Cuing Condition - Zero Disparity Target Cued | -0.055 | 0.346 | -0.160 | 0.873 | |
| Contrast : Cuing Condition - Uncued | -0.389 | 0.299 | -1.302 | 0.193 | |

| **B)** | **Estimates** | **2.5%** | **97.5%** |
|---|---|---|---|
| Intercept - Zero Disparity Distractor Cued | -0.650 | -1.262 | -0.041 |
| Cuing Condition - Crossed Disparity Target Cued | -0.073 | -0.400 | 0.268 |
| Cuing Condition - Crossed Disparity Distractor Cued | -0.239 | -0.588 | 0.130 |
| Cuing Condition - Uncrossed Disparity Target Cued | 0.007 | -0.364 | 0.384 |
| Cuing Condition - Uncrossed Disparity Distractor Cued | -0.247 | -0.599 | 0.116 |
| Cuing Condition - Zero Disparity Target Cued | 0.006 | -0.385 | 0.368 |
| Cuing Condition - Uncued | 0.195 | -0.094 | 0.523 |
| Contrast - Zero Disparity Distractor Cued | -2.212 | -2.674 | -1.744 |
| Contrast : Cuing Condition - Crossed Disparity Target Cued | -0.395 | -1.153 | 0.296 |
| Contrast : Cuing Condition - Crossed Disparity Distractor Cued | 0.498 | -0.173 | 1.146 |
| Contrast : Cuing Condition - Uncrossed Disparity Target Cued | -0.031 | -0.776 | 0.656 |
| Contrast : Cuing Condition - Uncrossed Disparity Distractor Cued | 0.722 | 0.091 | 1.353 |
| Contrast : Cuing Condition - Zero Disparity Target Cued | -0.055 | -0.805 | 0.629 |
| Contrast : Cuing Condition - Uncued | -0.389 | -0.955 | 0.189 |

**Table S3: Summary of the effects of the cuing conditions compared to the Uncued condition for trials without a distractor. A)** Estimates, standard errors, Wald test results and p-values for each factors in the selected model**. B)** Bootstrapped 95% confidence intervals of the estimates presented in table A).

| A) | Estimate | Std Error | Z Value | P-value | |
|---|---|---|---|---|---|
| Intercept - Uncued | -0.149 | 0.376 | -0.396 | 0.692 | |
| Cuing Condition - Crossed Disparity Target Cued | -0.199 | 0.195 | -1.019 | 0.308 | |
| Cuing Condition - Crossed Disparity Distractor Cued | -0.575 | 0.200 | -2.869 | 0.004 | ** |
| Cuing Condition - Uncrossed Disparity Target Cued | -0.362 | 0.198 | -1.830 | 0.067 | |
| Cuing Condition - Uncrossed Disparity Distractor Cued | -0.556 | 0.200 | -2.775 | 0.006 | ** |
| Cuing Condition - Zero Disparity Target Cued | -0.069 | 0.197 | -0.350 | 0.726 | |
| Cuing Condition - Zero Disparity Distractor Cued | -0.257 | 0.198 | -1.297 | 0.195 | |

| B) | Estimates | 2.5% | 97.5% |
|---|---|---|---|
| Intercept - Uncued | -0.149 | -0.867 | 0.645 |
| Cuing Condition - Crossed Disparity Target Cued | -0.199 | -0.548 | 0.180 |
| Cuing Condition - Crossed Disparity Distractor Cued | -0.575 | -0.996 | -0.179 |
| Cuing Condition - Uncrossed Disparity Target Cued | -0.362 | -0.776 | -0.006 |
| Cuing Condition - Uncrossed Disparity Distractor Cued | -0.556 | -0.959 | -0.116 |
| Cuing Condition - Zero Disparity Target Cued | -0.069 | -0.487 | 0.304 |
| Cuing Condition - Zero Disparity Distractor Cued | -0.257 | -0.633 | 0.148 |

**Table S4: Summary of effects of the cuing conditions compared to the Zero Disparity, Distractor cued condition for trials without a distractor. A)** Estimates, standard errors, Wald test results and p-values of factors in the selected model**. B)** Bootstrapped 95% confidence intervals of the estimates presented in table A).

| **A)** | **Estimate** | **Std Error** | **Z Value** | **P-value** |
|---|---|---|---|---|
| Intercept - Zero Disparity Distractor Cued | -0.405 | 0.394 | -1.029 | 0.303 |
| Cuing Condition - Crossed Disparity Target Cued | 0.058 | 0.227 | 0.256 | 0.798 |
| Cuing Condition - Crossed Disparity Distractor Cued | -0.318 | 0.232 | -1.375 | 0.169 |
| Cuing Condition - Uncrossed Disparity Target Cued | -0.106 | 0.230 | -0.460 | 0.646 |
| Cuing Condition - Uncrossed Disparity Distractor Cued | -0.299 | 0.231 | -1.293 | 0.196 |
| Cuing Condition - Zero Disparity Target Cued | 0.188 | 0.229 | 0.818 | 0.413 |
| Cuing Condition - Uncued | 0.257 | 0.198 | 1.297 | 0.195 |

| **B)** | **Estimates** | **2.5%** | **97.5%** |
|---|---|---|---|
| Intercept - Zero Disparity Distractor Cued | -0.405 | -1.186 | 0.309 |
| Cuing Condition - Crossed Disparity Target Cued | 0.058 | -0.376 | 0.506 |
| Cuing Condition - Crossed Disparity Distractor Cued | -0.318 | -0.797 | 0.153 |
| Cuing Condition - Uncrossed Disparity Target Cued | -0.106 | -0.562 | 0.386 |
| Cuing Condition - Uncrossed Disparity Distractor Cued | -0.299 | -0.765 | 0.218 |
| Cuing Condition - Zero Disparity Target Cued | 0.188 | -0.310 | 0.644 |
| Cuing Condition - Uncued | 0.257 | -0.132 | 0.657 |

**Table S5: Summary of the model investigating the effect of cue position for trials where the distractor is absent. A)** Estimates, standard errors, Wald test results and p-values of factors in the model**. B)** Bootstrapped 95% confidence intervals of the estimates presented in table A).

| A) | Estimates | Std Error | Z Value | P-value | |
|---|---|---|---|---|---|
| Intercept | -0.307 | 0.379 | -0.810 | 0.418 | |
| Distractor Cued | -0.269 | 0.136 | -1.975 | 0.048 | * |

| B) | Estimates | 2.5% | 97.5% |
|---|---|---|---|
| Intercept | -0.307 | -1.025 | 0.432 |
| Distractor Cued | -0.269 | -0.563 | -0.0001 |

**Figure S1: The influence of cue position and disparity on distractor saccade probability**. Mean (± 95% C.I.) proportion of trials with a first saccade towards the distractor as a function of its Michelson contrast (N=24 mantises). The cue was presented either on the side of the target (A) or on the side of the distractor (B). Blue curves represent trials with crossed disparity cues, yellow curves are for trials with uncrossed disparity cues while pink cues are for trial conducted with zero disparity cues. Green curves represent uncued trials and are the same across the two panels. The presented data combines the data from both 4 s (Experiment 1) and 100 ms (Experiment 2) cues. Overall, only the contrast of the distractor had a significant effect on the probability of first saccades towards the distractor (Model with contrast compared to null model: $\chi^2$=334.75, df=1, p<0.001 ; Estimate ± standard error of the contrast=2.440±0.145, Z=16.79, p<0.001).

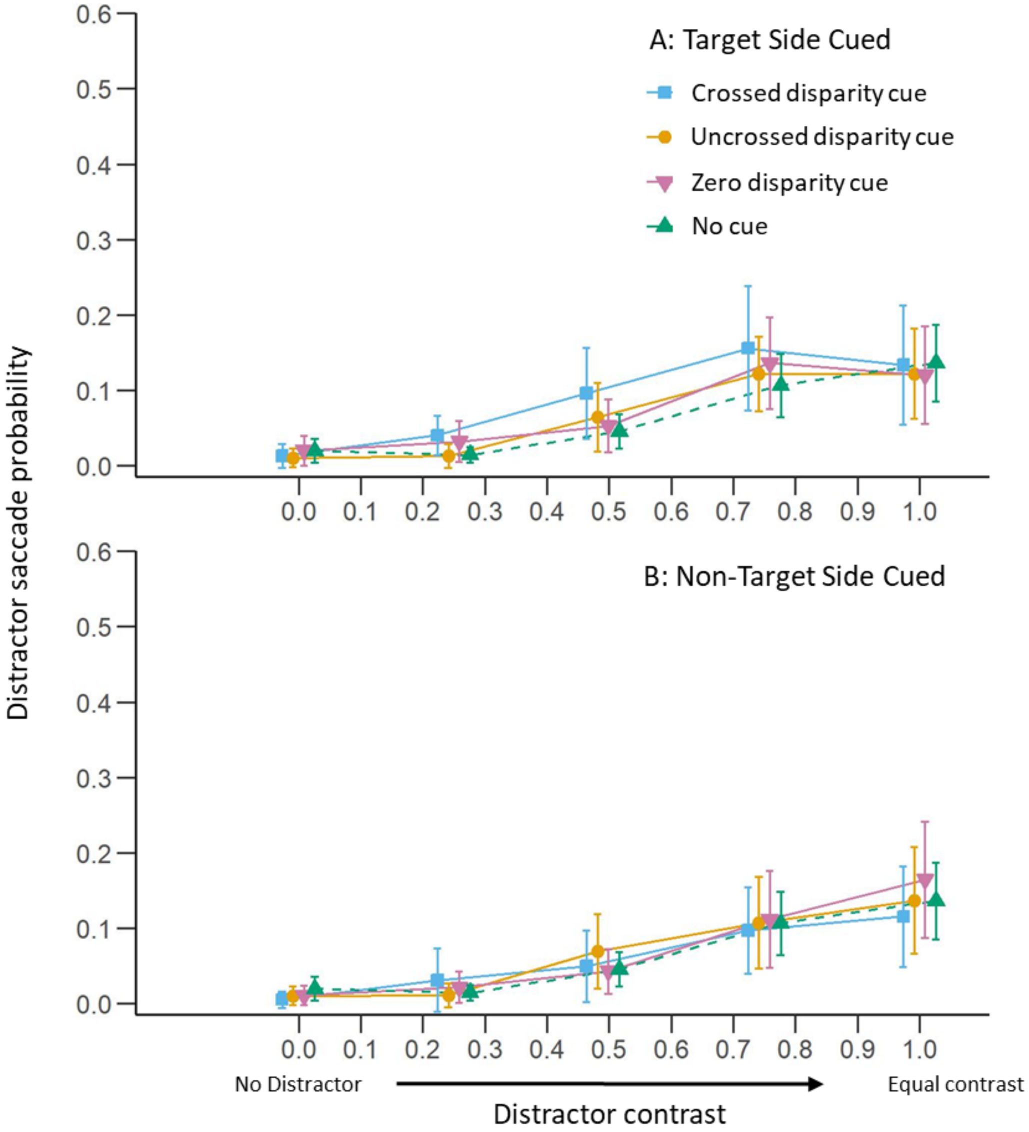

**Movie 1: Examples of trials presented to mantises during the experiment 1.** This experiment used 4 s cues. The first trial shows a crossed disparity cue placed on the target side and a distractor with a 0.75 contrast. The second trial shows an uncrossed disparity cue, on the target side and a 0.49 contrast distractor. The third trial shows a Zero disparity cue on the side opposite to the target and a 0 contrast (invisible) distractor. Finally, the fourth trial shows an Uncued condition with a 0.25 distractor contrast. Each trial is preceded by the centring stimulus. These trials were shown to a mantis placed upside-down in front of the screen with a purple filter in front of its left eye and the red filter in front of its right eye. Please note that crossed and uncrossed disparity cues and the target and distractor each consist of two images (one perceived through the purple filter and the other through the red one). However, to balance perception of the two images through the two filters, the gain of the image for the purple filter was strongly reduced. Thus, this image may be difficult to perceive depending on the reader's screen settings.

**Movie 2: Examples of trials presented to mantises during the experiment 2.** This experiment used 100 ms cues. The first trial presents a crossed disparity cue on the side of a full contrast the distractor. The second trial shows an uncrossed disparity cue presented on the target side and with a 0.25 contrast distractor. The third trial shows a Zero disparity cue placed on the distractor side, with a 0.49 contrast distractor. The fourth trial is Uncued with a 0.25 distractor. Please see additional remarks in the movie 1 caption.

**Movie 3: Examples of trials presented to mantises during the experiment 3.** In this experiment, the cue lasted 4 s and was always presented with a Zero disparity. Compared to the experiment 1 and 2, the cue could either be shifted towards the centre of the screen (centred) or towards the edge of the screen (excentred). The first trial has the cue centred and shown on the side of a 0.25 contrast distractor. The second trial also presents the cue on the side of the distractor but with the excentred condition and 0.75 distractor contrast. The third trial shows an excentred cue placed on the target side and with a 0 contrast distractor (invisible). Finally, the fourth trial shows a centred cue placed on the target side with a 0.49 contrast distractor. Please see additional remarks in the movie 1 caption.
